# Comparative analysis of soybean transcriptional profiles reveals defense mechanisms involved in resistance against *Diaporthe caulivora*

**DOI:** 10.1101/2023.03.27.534358

**Authors:** Eilyn Mena, Guillermo Reboledo, Silvina Stewart, Marcos Montesano, Inés Ponce de León

**Affiliations:** Departamento de Biología Molecular, Instituto de Investigaciones Biológicas Clemente Estable, Montevideo, Uruguay; Instituto Nacional de Investigación Agropecuaria, (INIA), Programa Nacional de Cultivos de Secano, La Estanzuela, Colonia, Uruguay; Laboratorio de Fisiología Vegetal, Centro de Investigaciones Nucleares, Facultad de Ciencias, Universidad de la República, Montevideo, Uruguay

**Keywords:** soybean stem canker, *Diaporthe caulivora*, transcriptomes, plant resistance, plant defense genes

## Abstract

Soybean stem canker (SSC) caused by the fungal pathogen *Diaporthe caulivora* is an important disease affecting soybean production. However, limited information related to the molecular mechanisms underlying soybean resistance to *Diaporthe* species is available. In the present work, the defense responses to *D. caulivora* in two contrasting soybean genotypes were analyzed. We showed that *Génesis* 5601 is more resistant to fungal infection than Williams, evidenced by significantly smaller lesion length, reduced disease severity and pathogen biomass. Transcriptional profiling was performed in untreated plants and in *D. caulivora*-inoculated and control-treated tissues at 8 and 48 hours post inoculation (hpi). In total, 2.322 and 1.855 genes were differentially expressed in Génesis 5601 and Williams, respectively. Interestingly, Génesis 5601 exhibited a significantly higher number of upregulated genes compared to Williams at 8 hpi, 1.028 versus 434 genes. Resistance to *D. caulivora* was associated with defense activation through transcriptional reprogramming mediating perception of the pathogen by receptors, biosynthesis of phenylpropanoids, hormones, small heat shock proteins and pathogenesis related (PR) genes. These findings provide novel insights into soybean defense molecular mechanisms used to control *D. caulivora*, and generate a foundation for development of resistant SSC varieties within soybean breeding programs.

## Introduction

Soybean (*Glycine max* L.) is a major global crop affected by biotic stress caused by microbial pathogens, nematodes and insects, as well as abiotic stress such as drought, nutrient deficiency, salt and cold^1^. Soybean stem canker (SSC), caused by fungal *Diaporthe* species is an important soybean disease worldwide. *D. aspalathi, D. caulivora* and *D. longicolla* are the principal agents causing SSC in different countries^2-5^. Disease symptoms appear on the stem as 1-2 mm spots that expand as elongated brown lesions associated to withered brown leaves^3^. SSC control is based on integrating management practices such as crop rotation and fungicides application. However, the most effective way to control SSC is to develop and use resistant cultivars. Five *Rdm* loci confer resistance to *D. aspalathi*^3,6^, although these resistant genes are not effective against *D. caulivora*^3^. Recently, an *Rdc1* locus of *G. max* was identified as a resistance source for *D. caulivora*^7^. However, the molecular identity of Rdc1 is currently unknown.

Plants perceive pathogens and trigger cellular and molecular modifications associated with defense responses such as signaling, transcriptional activation, synthesis of defense molecules and their transport to specific sites in the plant^8^. Recognition occurs at the plasma membrane by pattern-recognition receptors (PRRs) and in the cytoplasm by nucleotide-binding domain leucine-rich repeat containing receptors (NLRs)^9,10^. PRRs perceive conserved pathogen-or damage-associated molecular patterns (PAMPs or DAMPs) at the surface of the plant cells to activate pattern-triggered immunity (PTI). PRRs include receptor-like kinases (RLKs) or receptor-like proteins (RLP) with different extracellular domains. NLRs perceive pathogen effectors delivered inside the plant cell leading to effector-triggered immunity (ETI). Both PTI and ETI activate overlapping events such as mitogen-activated protein kinases (MAPKs) cascades, Ca2+ flux, hormonal signaling, reactive oxygen species (ROS) burst, callose deposition, and transcriptional reprogramming^11,12^.

During soybean-*D. caulivora* interaction, plant cells activate the expression of genes encoding pathogenesis-related proteins (PR-1, PR-2, PR-3, PR-4, PR-10), and enzymes involved in phenylpropanoid and oxylipin synthesis (PAL, phenylalanine-ammonia lyase; CHS, chalcone synthase and LOX, lipoxygenases)^2^. Most of these defense genes were also induced in soybean tissues infected with *D. aspalathi*^13^. The phenylpropanoid and oxylipin pathways participate in plant defense against pathogens by producing important compounds with antimicrobial activities, and contribute to reinforcement of the cell wall and defense signaling^14,15^. Recently, we sequenced the genome of *D. caulivora* (isolate D57) and performed transcriptomic analysis during stem colonization to reveal the molecular basis of fungal pathogenesis^16^. *D. caulivora* infection strategy relies on plant cell wall degradation and modification, detoxification of plant compounds, transporter activities, toxin production and effector proteins involved in plant defense evasion^16^. However, the molecular mechanisms employed by plant cells to recognize *D. caulivora* and activate an effective defense response leading to resistance are still unknown.

In other soybean-pathogen interactions, transcriptomic studies have allowed to understand complex gene regulatory networks operating in plants infected with different pathogens, including *Phytophthora sojae*^17^, *Phakopsora pachyrhizi*^18,19^, *Fusarium oxysporum*^20^, and soybean mosaic virus^21^. To unravel the mechanisms involved in defense activation and resistance against *D. caulivora*, we performed RNAseq profiling in two contrasting soybean genotypes, the susceptible cultivar Williams and the resistant cultivar Génesis 5601. The results revealed a complex and differential gene expression network during the activation of defense responses between cultivars upon pathogen infection.

## Results

### *D. caulivora* infection of soybean genotypes

Williams and Génesis 5601 cultivars were inoculated with *D. caulivora* and SSC progress was monitored during 14 days post inoculation (dpi). Disease symptoms were clearly visible in Williams, while Génesis 5601 exhibited only small stem lesions (Figure 1a). Lesions expanded in both directions of the stem and only in Williams we observed withered leaves above de canker lesion. From 5 dpi, lesion length was significantly higher in Williams compared to Génesis 5601, varying between 36 to 50% (Figure 1a,b). Disease severity index and area under the disease progress curve (AUDPC) were evaluated in soybean stems until 14 dpi. According to a disease severity scale [2], *D. caulivora* infection resulted in more symptom development in Williams compared to Génesis 5601 throughout time (Figure 1c-d). In addition, *D. caulivora* biomass was quantified in soybean stems of both genotypes by quantitative PCR (qPCR). Fungal DNA (ng *D. caulivora*/ng soybean) was measured at 8, 24, 48, 72 and 96 hours post inoculation (hpi). The results show that pathogen biomass was significantly higher in Williams compared to Génesis 5601 at 24-96 hpi (2-4 fold) (Figure 1e).

**Figure 1.**
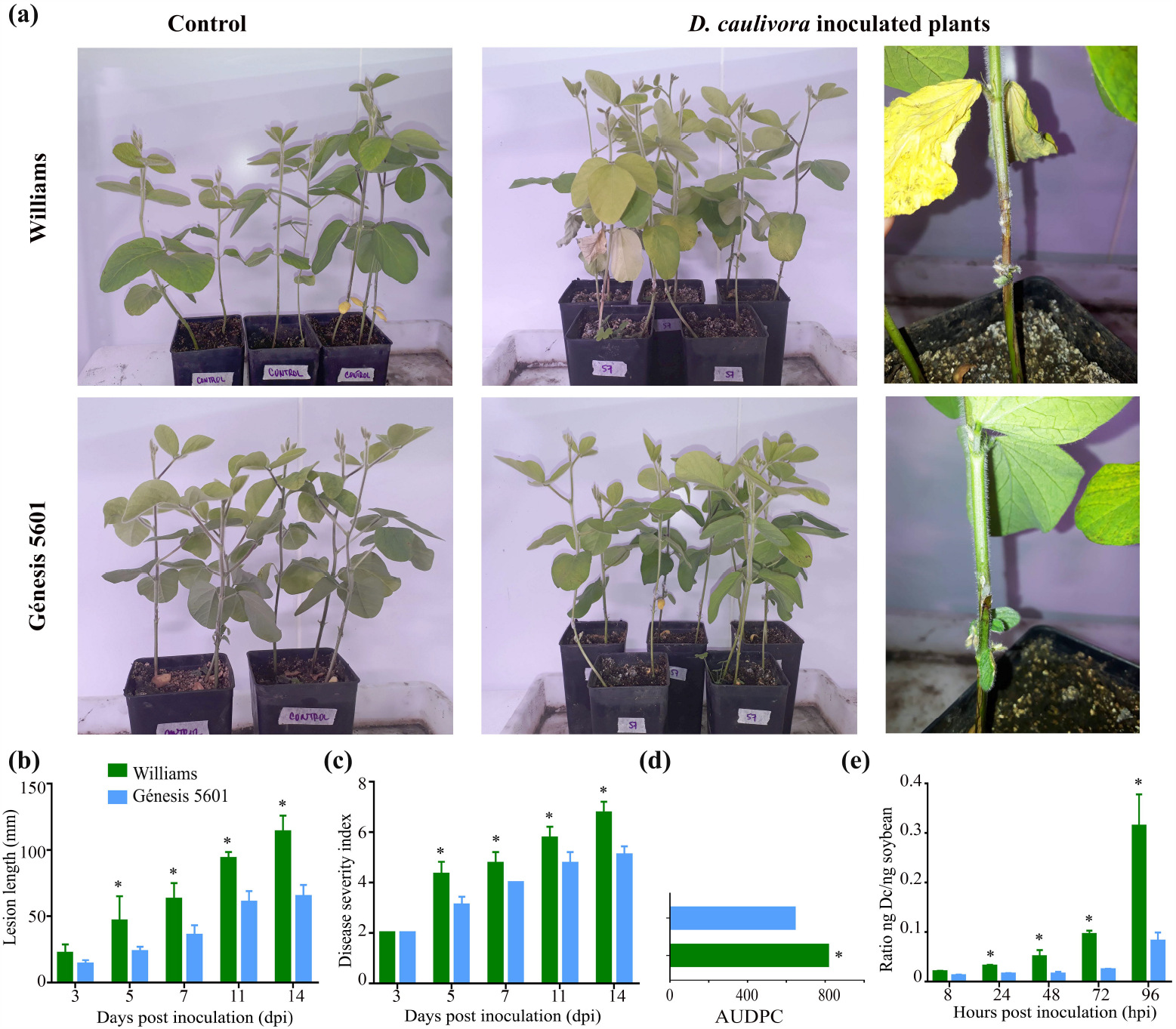
Soybean stem canker disease progress after *D. caulivora* inoculation. **(a)** Symptoms in susceptible and resistant soybean plants following *D. caulivora* inoculation at 7 days post-inoculation (dpi), **(b)** Lesion length in susceptible and resistant soybeans at 3, 5, 7, 11 and 14 dpi, **(c)** Disease severity index in susceptible and resistant soybeans at 3, 5, 7, 11 and 14 dpi, **(d)** AUDPC in susceptible and resistant soybeans at 3, 5, 7, 11 and 14 dpi. *D. caulivora* biomass in susceptible and resistant soybeans at 8, 24, 48, 72 and 96 hpi. *Indicates a significant difference between the soybean genotypes at p-value <0.05 (One-way ANOVA).

### Transcriptome profiles of contrasting soybean genotypes infected with *D. caulivora*

To identify molecular components involved in soybean defense against *D. caulivora*, we first compared the transcriptomes of Williams and Génesis 5601 under normal conditions in untreated plants. A second comparison included Williams and Génesis 5601 tissues inoculated with *D. caulivora* versus control treatment at the indicated time points. Soybean stem samples were taken from untreated plants, and stem tissues from *D. caulivora*-inoculated and control-treated plants were collected at 8 and 48 hpi. A total of 737 million reads were generated after removing adapter sequences and low-quality reads. Reads mapped uniquely to *G. max* nuclear genome were considered for further analyses (18.2-34.5 million reads), with 83-98% of reads mapping to the reference genome of soybean (Supplementary Table S1). The principal component analysis of the different treatments showed a clear separation, and variability among biological replicates was very low as indicated (Supplementary Figure S1).

Soybean expression profiles were analyzed in Génesis 5601 and Williams during *D. caulivora* inoculation at 8 and 48 hpi compared to their respective controls (from now on Génesis-8 and -48 and Williams-8 and -48). The analysis revealed a total of 2.384 differentially expressed genes (DEGs) between soybean genotypes (Supplementary Table S2). A massive transcriptional shift towards upregulation was observed at 8 and 48 hpi after *D. caulivora* inoculation compared to control treatment (Figure 2a). Untreated Génesis 5601 plants exhibited 164 DEGs (73 up- and 91 downregulated) compared to untreated Williams plants. During infection, Williams-48 has more DEGs than Williams-8, 1.342 and 513, respectively (Figure 2a, Supplementary Table S2). However, the number of DEGs in Génesis-8 and Génesis-48 were similar, 1.115 and 1.207, respectively, indicating that Génesis-8 has significantly more DEGs than Williams-8. Among DEGs, 167 were upregulated in both genotypes at both time points, while some genes were uniquely expressed in one condition (Figure 2b). A total of 930 DEGs were commonly upregulated in both genotypes, representing 46 % of the upregulated DEGs (2.011 genes). In contrast, 22 DEGs were downregulated in both genotypes, representing 5,9 % of a total of 376 genes. Furthermore, 75 and 327 upregulated DEGs were only found in Williams-8 and Williams-48, and 401 and 191 upregulated DEGs were only present in Génesis-8 and Génesis-48, respectively (Figure 2b).

**Figure 2.**
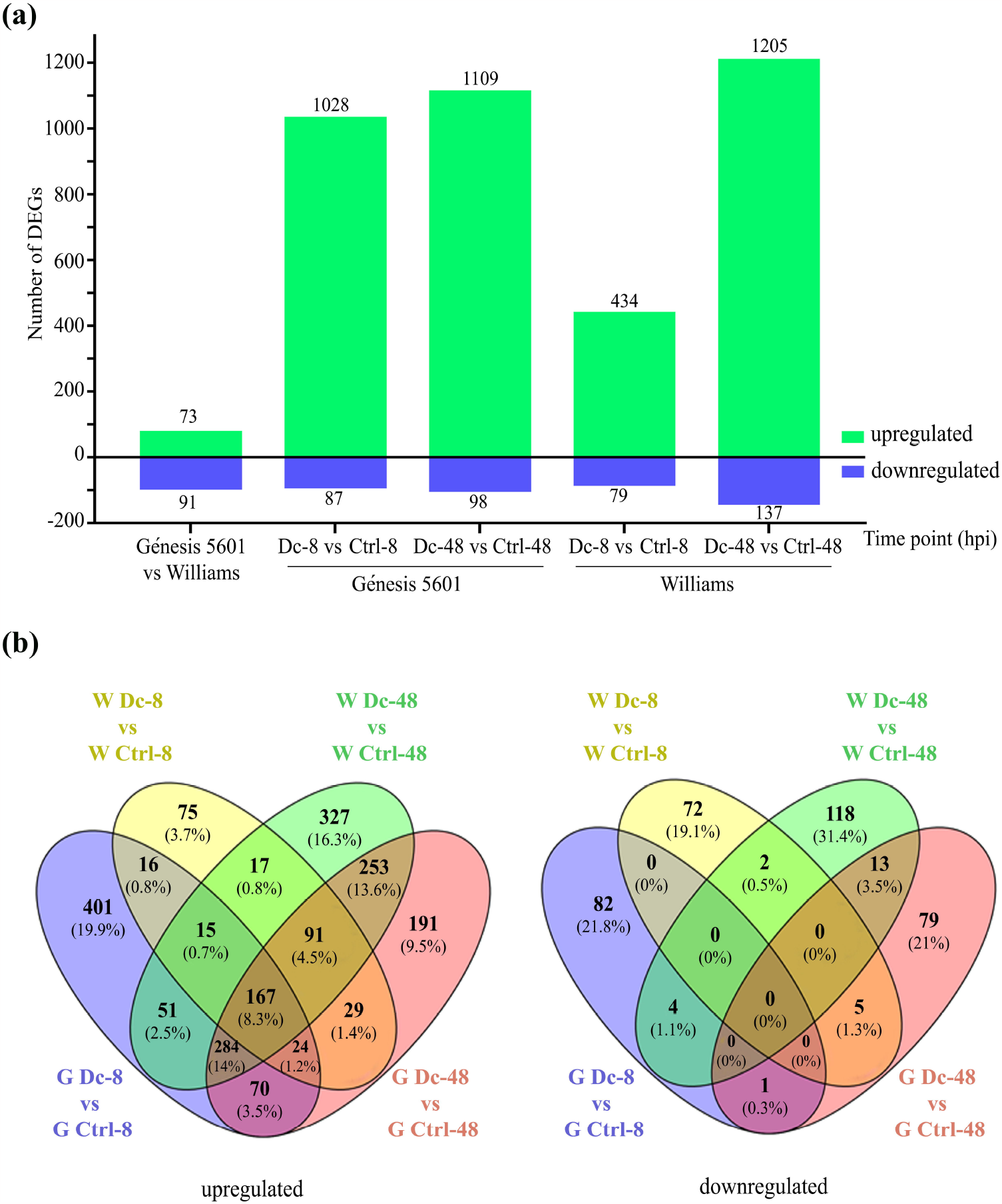
Differentially expressed genes (DEGs) identification in susceptible (Williams) and resistant (Génesis 5601) soybean plants without treatment and after *D. caulivora* inoculation. **(a)** Number of DEGs for each treatment in both genotypes. Log2 FC≥2.0 or ≤2.0 and false discovery rate (FDR) ≤ 0.05 were considered for DEGs identification, **(b)** Venn diagram showing the number of upregulated and downregulated soybean genes at 8 and 48 hours post inoculation (hpi) with *D. caulivora* in susceptible and resistant soybean plants. Overlap of expressed soybean genes can be observed.

### Functional enrichment of differentially expressed genes

We further focused on upregulated genes in the different comparisons by analyzing Gene Ontology (GO) term enrichment analysis and manual inspection in order to identify biological processes (BP) and molecular functions (MF). Enriched GO were not found for untreated Williams or Génesis 5601 tissues. Most of the top 25 significantly enriched GO terms in *D. caulivora* inoculated versus control soybean plants were similar at 8 and 48 hpi (Figure 3). Génesis 5601 showed a greater number of genes per category respect to Williams principally at 8 hpi. The most represented BP in both genotypes at 8 and 48 hpi were protein phosphorylation, regulation of transcription and defense. Other top GO terms enrichment in BP at 8 hpi, included ethylene-activated signaling pathway, response to oxidative stress and hydrogen peroxide transmembrane transport, response to heat and salt stress, ABA activated signaling pathway, response to auxin and cell wall modification. All these GOs related to defense were also present at 48 hpi and additional upregulated defense-related GO terms included cellular oxidant detoxification, flavonoid biosynthetic process, and response to biotic stimulus, bacterium, chitin and fungus. In general, biotic related process, abiotic related process, hormones and secondary metabolites represent 60 % of the GO biological process (BP) terms identified in the up-regulated genes. In both genotypes the MF terms are mostly represented by ATP binding and DNA-binding transcription factor activity at 8 and 48 hpi. Top MF terms included ATP-, heme-, protein-, DNA- and iron ion-binding, sequence-specific DNA binding, protein kinase-, oxidoreductase-, monooxygenase-, peroxidase-, glycosyltransferase- and protein serine threonine kinase-activity.

**Figure 3.**
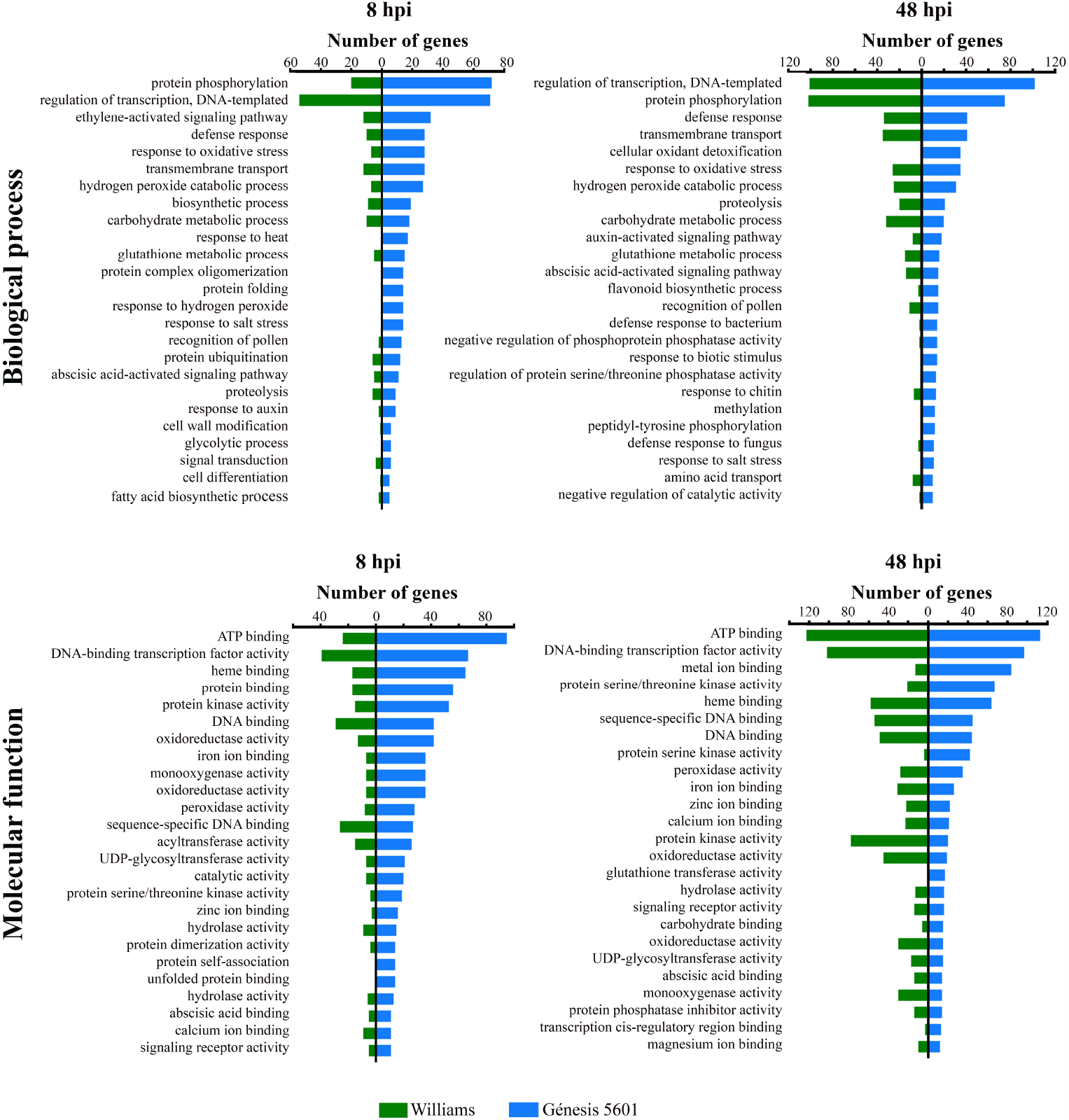
Enriched gene ontology (GO) biological process and molecular function terms. Top 25 enrichment GO terms (p<0.05) of upregulated genes in Williams (green bars) and Génesis 5601 (blue bars) at 8 and 48 hours post inoculation (hpi) with *D. caulivora*.

To study the host metabolic pathways altered during *D. caulivora* infection, we performed a KEGG enrichment analysis. The majority of enriched KEGG pathways were related to plant defense and plant-pathogen interaction, including phenylpropanoid, flavonoid, flavone and flavonol, isoflavonoid and anthocyanin biosynthesis, plant hormone signal transduction, brassinosteroid and steroid hormone biosynthesis. Both genotypes shared most of the KEGG pathways, although there were more genes and unique pathways associated to Génesis 5601 than to Williams. The most enriched KEGG pathways for upregulated DEGs in Génesis 5601 were secondary metabolites biosynthesis, plant hormone signal transduction and plant-pathogen interaction.

### Differential expression of genes involved in plant defense during *D. caulivora* infection

Hierarchical clustering was performed to group similar expression patterns across genotypes and treatments. This analysis grouped Williams-48 with Génesis-48, and Génesis-8 was more related to this group than to Williams-8, which is consistent with a significantly higher number of upregulated DEGs in Génesis-8 (Figure 4, Supplementary Table S3). The analysis of the total DEGs identified six clusters with different expression patterns. To analyze the resistant mechanisms of soybean plants against *D. caulivora* infection, we further focused on cluster 4, 5, and 6. Cluster 4 contained 400 DEGs that were only upregulated in Génesis-8, including genes involved in functional categories such as perception of the pathogen, hormonal signaling, transcription factors (TFs), phenylpropanoid biosynthesis, transporters, ROS detoxification and PR genes. Cluster 5 comprises 191 DEGs that were only upreglated in Génesis-48 and 24 DEGs that were upregulated in Génesis-8 and Génesis-48. Cluster 6 contains 255 DEGs that were only upregulated in Génesis-48 and Williams-48 and 280 DEGs that were upregulated in Génesis-8, Génesis-48 and Williams-48. Cluster 5 and 6 have genes in the same functional categories as Cluster 4.

**Figure 4.**
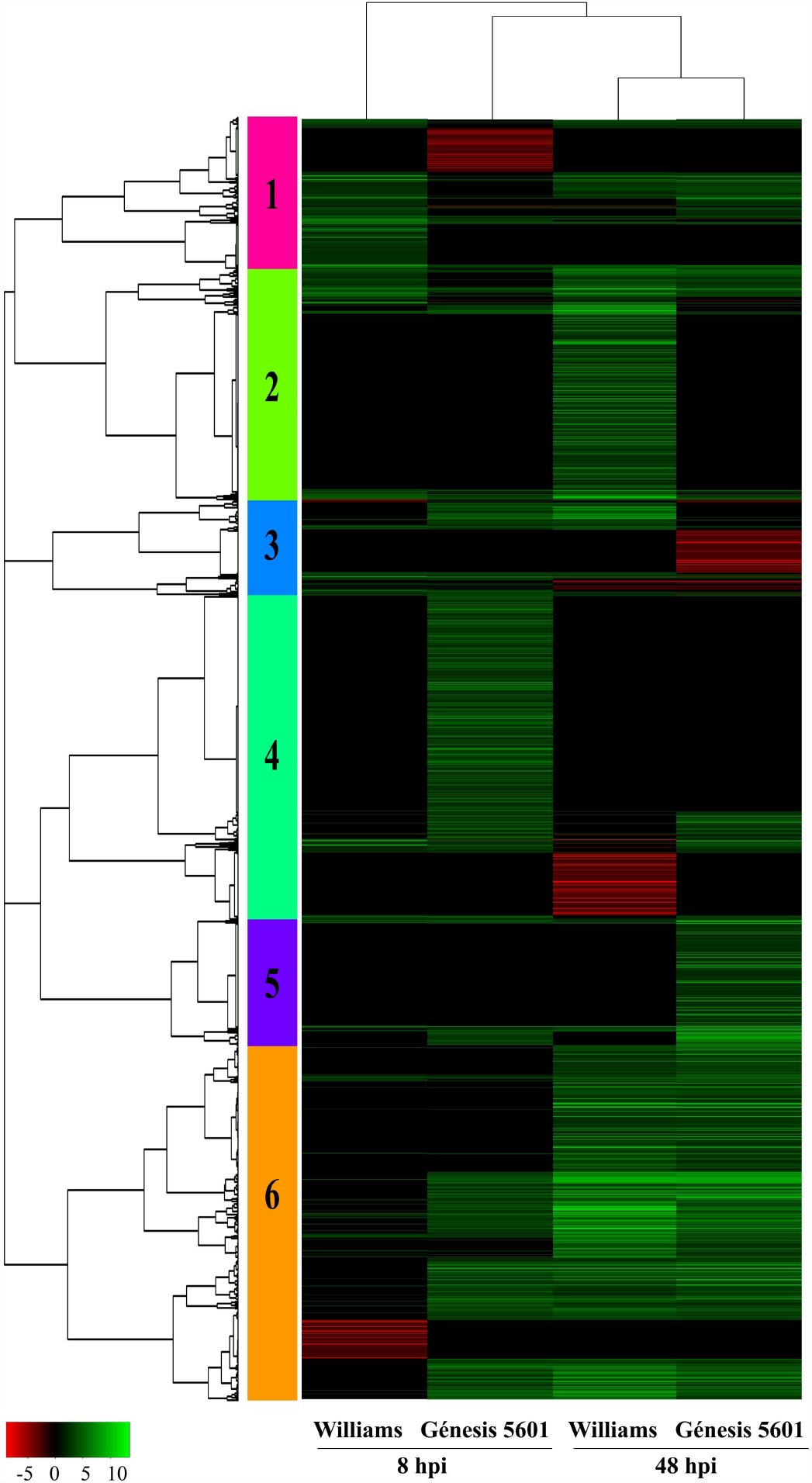
Heat map of hierarchical clustering showing all differentially expressed genes (DEGs) in soybean plants inoculated with *D. caulivora*. Green (upregulated) and Red (downregulated). See Supplementary table S3 for complete information.

By performing a deeper inspection of genes within functional categories among genotypes, we found that most of them were upregulated in both genotypes, although as previously mentioned, the number of genes was significantly higher in Génesis-8 compared to Williams-8. However, we found a group of DEGs that encodes small heat shock proteins (sHSPs) that were only upregulated in Génesis 5601, while they did not show differential expression in Williams. In total, 17 sHSPs and one sHSP were upregulated at Génesis-8 and Génesis-48, respectively. Moreover, three Bcl-2-associated athanogene (BAG) cochaperone encoding genes, Glyma.18G284900, Glyma.18G285100 and Glyma.07G061500, were only upregulated in Génesis 5601.

### Induced expression of genes involved in pathogen perception, signaling and transcription during soybean infection by *D. caulivora*

Pathogen recognition, signaling and transcriptional reprogramming are important steps in the activation of plant defense mechanism against pathogens. We identified 159 DEGs encoding PRRs, most of which were upregulated during *D. caulivora* infection, including leucine-rich repeat receptor-like protein kinase (LRR-RLK), RLKs, RLPs and lectin domain containing receptor kinase (LecRLKs), among others (Figure 5, Supplementary Table S4). While only 11 receptor genes were upregulated in Williams-8, the number of these upregulated genes increased to 59 in Génesis-8, including LRR-RLKs, RLPs, cysteine-rich receptor-like protein kinase (CRKs) and LecRLKs. The number of upregulated receptor genes were 76 in Génesis-48 and 94 in Williams-48. Furthermore, 15 upregulated DEGs encoded NLR; seven in Génesis-8, one in Williams-8, three in Génesis-48 and eight in Williams-48. In untreated plants, a higher number of PRRs and NLR genes showed increased expression levels in Génesis 5601 compared to Williams.

**Figure 5.**
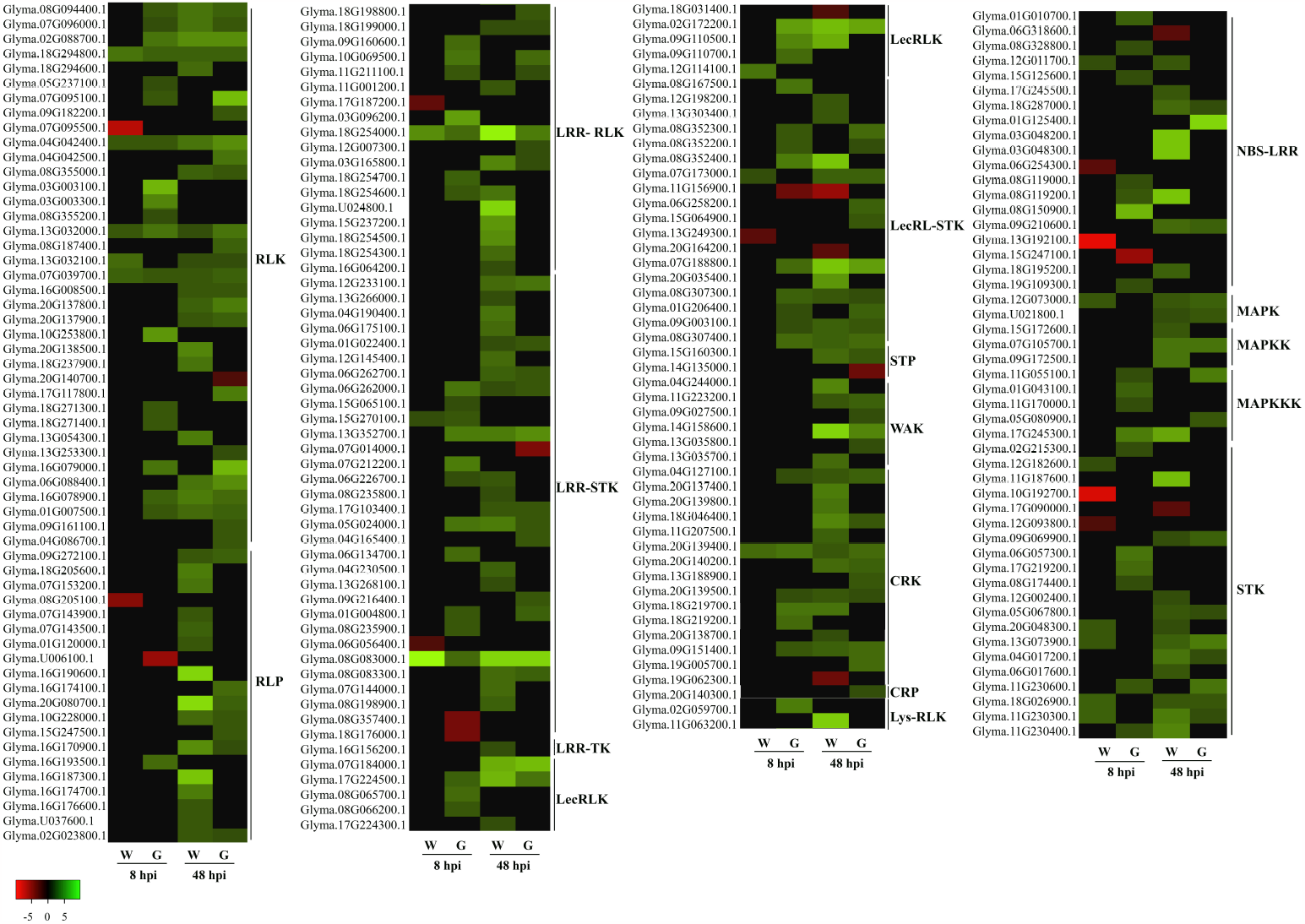
Heat map of differentially expressed genes (DEGs) encoding for proteins with roles in perception and signaling. Individual genes are listed and colors represented the log2 fold change value based on the comparison of the transcript levels between *D. caulivora* inoculated and control treatment for both genotypes (Williams and Génesis 5601). Green (upregulated) and Red (downregulated). Abbreviated are as follows: Receptor-like protein kinase (RLK), Receptor like protein (RLP), Leucine-rich repeat receptor-like protein kinase (LRR-RLK), Lectin domain containing receptor kinase (LecRLK), wall-associated receptor kinase (WAK), Cysteine-rich receptor-like protein kinase (CRK), Cysteine-rich protein (CRP), LysM domain receptor-like kinase (LysM-RLK), Nucleotide-binding site leucine-rich repeat (NLR), Mitogen-activated protein kinase (MAPK), Mitogen-activated protein kinase kinase (MAPKK), Mitogen-activated protein kinase kinase kinase (MAPKKK), and Serine/threonine-protein kinase (STK).

Thirty DEGs encoded members of the mitogen-activated protein kinase (MAPK) cascade, including MAPK, MAPKK and MAPKKK and serine/threonine-protein kinase (STKs) (Figure 5, Supplementary Table S4). Four MAPKKK were only induced in Génesis-8 and different STKs were upregulated in Génesis-8 and Williams-8. Moreover, 219 DEGs encoded TFs related to plant defenses to biotic stress, including 48 WRKYs, 71 Apetala2/ethylene responsive factor (AP2/ERFs), 28 myeloblastosis related (MYB), 17 basic helix–loop–helix (bHLH), 15 no apical meristem, ATAF1/2, and cup-shaped cotyledon (NAC) and 8 Gibberellin-insensitive, repressor of GA1–3, and Scarecrow (GRAS) (Figure 6, Supplementary Table S5). Most TFs were upregulated during *D. caulivora* infection and the number of upregulated TFs increased significantly at 48 hpi respect to 8 hpi in both genotypes. Génesis-8 exhibited more upregulated *ERFs, MYBs* and *bHLH* compared to Williams-8.

**Figure 6.**
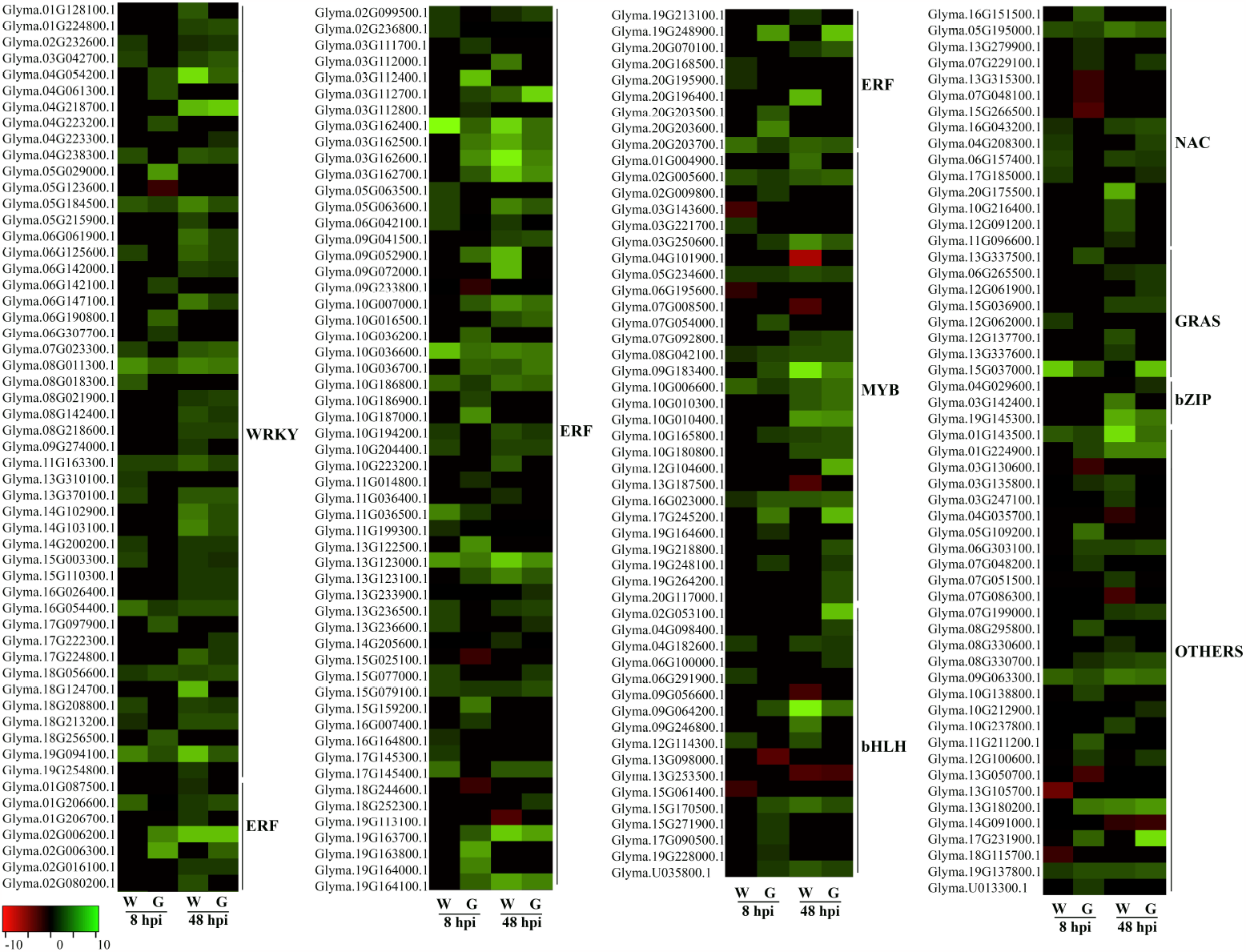
Heatmap of differentially expressed genes (DEGs) encoding for transcription factors. Individual genes are listed and colors represented the log2 fold change value based on the comparison of the transcript levels between *D. caulivora* inoculated and control treatment for both genotypes (Williams and Génesis 5601). Green (upregulated) and Red (downregulated). Abbreviated are as follows: WRKY transcription factors (WRKY), Ethylene responsive transcription factor (ERF), MYB transcription factor (MYB), bHLH transcription factor (bHLH), Cup-shaped cotyledon (NAC), Gibberellin-insensitive, repressor of GA1-3 and Scarecrow (GRAS), bZIP transcription factor (bZIP), and others.

### Activation of pathogenesis-related genes and the phenylpropanoid pathway during *D. caulivora* infection

Pathogenesis-related (PR) proteins play important functions in plant immune responses^22^. During *D. caulivora* infection, soybean induced the expression of 139 PR genes encoding PR-1, β-1,3-glucanases (PR-2), chitinases (PR-3, PR-4, PR-8), thaumatins (PR-5), proteinase inhibitor (PR-6), endoproteinasse (PR-7), peroxidases (PR-9), PR-10 (ribonuclease-like protein), defensing (PR-12), lipid-transfer protein (PR-14) and germin-like proteins (PR-16) (Supplementary Table S6). A higher number of genes encoding PRs were upregulated in Génesis-8 compared to Williams-8, while at 48 hpi the number of upregulated genes were similar among genotypes. Four PR-2 were only induced in Génesis-8 and not in Williams-8, while six PR-14 were induced in Génesis-8 and only one in Williams-8. Similarly, 27 PR-9 and 10 PR-10 were upregulated in Génesis-8, while this number decreased to seven and four in Williams-8.

The phenylpropanoid pathway produces multiple compounds involved in defense mechanisms against biotic stress^14^. In total, 169 DEGs related to this pathway were identified during *D. caulivora* infection. A high number of genes encoding enzymes involved in flavonoids, isoflavonoids, flavonone, flavonols, flavones and anthocyanins biosynthesis were upregulated in both genotypes at 48 hpi (Figure 7, Supplementary Table 7). The most remarkable differences between genotypes were observed in Génesis-8, which has significantly more upregulated DEGs than Williams-8 (117 versus 50). For example, while 10 cytochrome P450 (CYP) and 14 CHS encoding genes were induced in Génesis-8, this number decreased to two *CYP* and eight *CHS* in Williams-8. Similarly, Génesis-8 showed increased expression of genes encoding for three isoflavone 7-O-methyltransferase (IOMT) and four isoflavone 2’-hydroxylase (*I2‥H*), while Williams-8 showed only increased expression of one *I2‥H* and differential expression of IOMT genes could not be observed. Other genes encoding dirigent proteins, involved in the synthesis of lignin, were also upregulated during *D. caulivora* infection in both genotypes.

**Figure 7.**
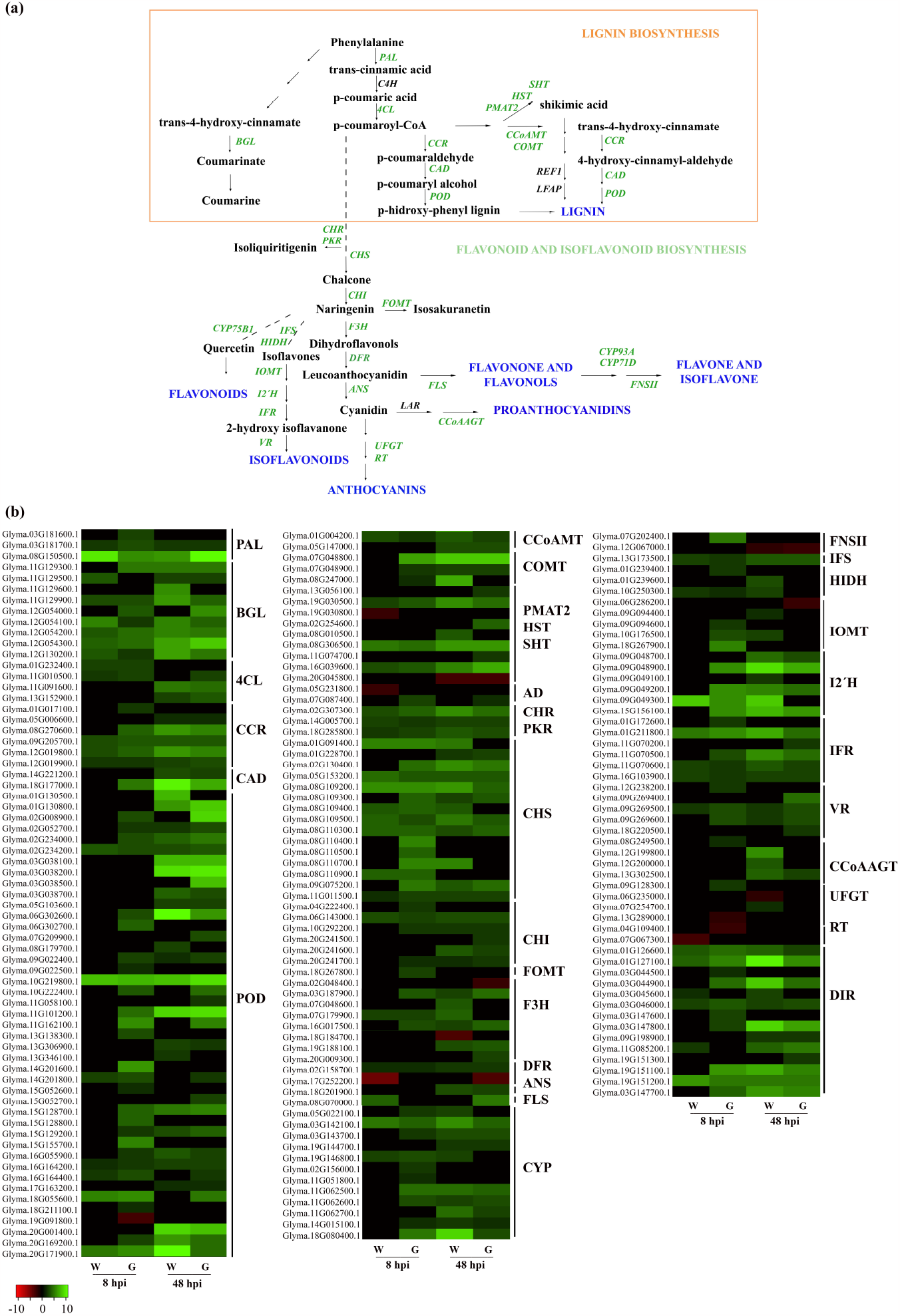
Activation of the phenylpropanoid pathway in response to *D. caulivora*. **(a)** Simplified scheme of phenylpropanoid and flavonoids biosynthetic pathway. Upregulated genes encoding enzymes of this pathway are highlighted in green. (b) Heatmap of differentially expressed genes (DEGs) encoding genes of the phenylpropanoid and flavonoids biosynthetic pathway. Individual genes are listed and colors represented the log2 fold change value based on the comparison of the transcript levels between *D. caulivora* inoculated and control treatment for both genotypes (Williams and Génesis 5601). Green (upregulated) and Red (downregulated). Abbreviated are as follows: Phenylalanine ammonia-lyase (PAL), beta-glucosidase (BGL), 4-coumarate--CoA ligase (4CL), cinnamoyl-CoA reductase (CCR), Cinnamyl alcohol dehydrogenase (CAD), Peroxidase (POD), Caffeoyl-CoA O-methyltransferase (CCoAMT), Caffeic acid 3-O-methyltransferase (COMT), Phenolic glucoside malonyltransferase 1-like (PMAT2), Spermidine hydroxycinnamoyl transferase (HST) Shikimate O-hydroxycinnamoyl transferase (SHT), Aldehyde dehydrogenase (AD), Chalcone reductase (CHR), Chalcone synthase (CHS), Chalcone isomerase (CHI), Isoflavone 7-O-methyltransferase (FOMT), Flavanone 3-hydroxylase (F3H), Dihydroflavonol-4-reductase (DFR), 2-oxoglutarate-dependent dioxygenase (ANS), DMR6-like oxygenase 2 (FLS), Cytochrome P450 (CYP), Flavone synthase II (FNSII), Isoflavone synthase 2 (IFS), 2-hydroxyisoflavanone dehydratase (HIDH), Isoflavone 7-O-methyltransferase (IOMT), Isoflavone 2’-hydroxylase (I2’H), Isoflavone reductase (IFR), Vestitone reductase (VR), Coumaroyl-CoA:anthocyanidin 3-O-glucoside-6’’-O-coumaroyltransferase (CCoAAGT), UDP-glycosyltransferase (UFGT), UDP-rhamnose:rhamnosyltransferase (RT), and Dirigent protein (DIR).

### Plant hormones involved in soybean defense against *D. caulivora*

GO enrichment analysis and overrepresented DEGs showed that salicylic acid (SA), auxin, ET, jasmonic acid (JA), abscisic acid (ABA), cytokinins (CK) and brassinosteroids (BR) probably participate in defense responses against *D. caulivora*. In total, 131 DEGs were involved in phytohormone pathways. Three PAL encoding genes with possible roles in SA synthesis were upregulated with *D. caulivora*, and two of these PAL genes were only upregulated in Génesis 5601. However, isochorismate synthase genes (ICSs) were not induced after *D. caulivora* inoculation (Figure 8, Supplementary Table S8). Auxin related DEGs included 37 genes involved in biosynthesis, signaling and response to indole-3-acetic acid (IAA), and most of them were upregulated. A higher number of small auxin-up RNA (SAUR) genes were induced in Génesis-8 and in Williams-48 (nine and 10), compared to two SAUR genes in Williams-8 and four in Génesis-48 (Figure 8, Supplementary Table S8). Furthermore, significantly more upregulated DEGs involved in CK synthesis and signaling were observed in Génesis-8 compared to Williams-8. We found three abscisate beta-glucosyltransferase, and three PYL4 ABA receptors that were upregulated with *D. caulivora*. Moreover, one ET receptor (ETR) was upregulated in both genotypes at 48 hpi, and 34 DEGs were related to ET synthesis and signaling; 24 were upregulated in Génesis-8 and only seven in Williams-8. Most of these DEGs were 1-aminocyclopropane-1-carboxylate synthase (ACC), 1-aminocyclopropane-1-carboxylate oxidase (ACO) and ERF TFs. Nine upregulated DEGs encoded proteins involved in JA biosynthesis and signaling in Génesis-8 such as LOX, 12-oxophytodienoate reductase (OPR) and MYC2 TF, while in Williams-8 only one OPR was upregulated. At 48 hpi, the number of DEGs related to ET and JA pathways were similar among soybean genotypes. Taken together, these results suggest that several hormones, especially IAA, ET, CK and JA are involved in resistance against *D. caulivora*.

**Figure 8.**
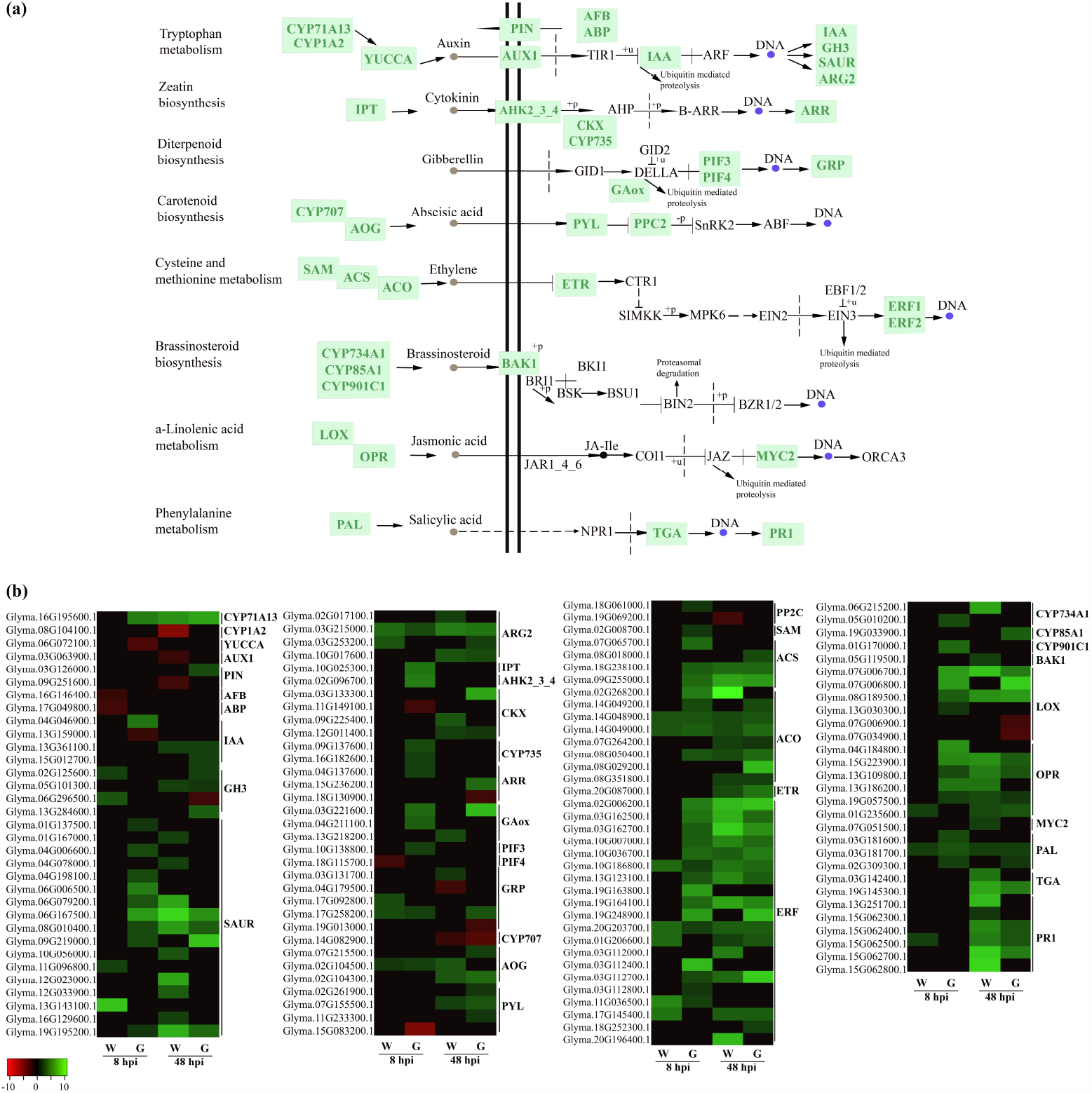
Activation of hormonal pathways after *D. caulivora* inoculation. **(a)** Simplified scheme of hormones signaling. Upregulated (green) genes are represented. (b) Heatmap of differentially expressed genes (DEGs) encoding enzymes of defense hormone signaling. Individual genes are listed and colors represented the log2 fold change value based on the comparison of the transcript levels between *D. caulivora* inoculated and control treatment for both genotypes (Williams and Génesis 5601). Green (upregulated) and Red (downregulated). Abbreviated are as follows: Cytochrome P450 (CYP71A13, CYP1A2), indole-3-pyruvate monooxygenase YUCCA (YUCCA), Auxin influx carrier (AUX1), Auxin efflux carrier (PIN), Auxin signaling F-box (AFB), Auxin-binding protein (ABP), Auxin responsive protein (IAA), Auxin responsive GH3 gene family (GH3), SAUR family protein (SAUR), Indole-3-acetic acid-induced protein ARG2 (ARG2), Adenylate isopentenyltransferase (IPT), Arabidopsis histidine kinase 2/3/4, a Cytokinin receptor (AHK2_3_4), Cytokinin dehydrogenase (CKX), Cytokinin hydroxylase (CYP735), Two-component response regulator ARR family (ARR), Gibberellin oxidase (GAox), Phytochrome-interactin factor 3 (PIF3), Phytochrome-interactin factor 4 (PIF4), Gibberellin-regulated protein (GRP), Abscisic acid 8’-hydroxylase (CYP707), Abscisate beta-glucosyltransferase (AOG), Abscisic acid receptor PYR/PYL family (PYL), Protein phosphatase 2C (PP2C), S-adenosylmethionine synthase (SAM), 1-aminocyclopropane-1-carboxylate synthase (ACS), 1-aminocyclopropane-1-carboxylate oxidase (ACO), Ethylene receptor (ETR), Ethylene-responsive transcription factor (ERF), Cytochrome P450 (CYP734A1, CYP85A1, CYP901C1), Brassinosteroid Insensitive 1-associated receptor 1 (BAK1), Linoleate lipoxygenase (LOX), 12-oxophytodienoate reductase (OPR), Transcription factor MYC2 (MYC2), Phenylalanine ammonia-lyase (PAL), Transcription factor TGA (TGA), Pathogenesis-related protein 1 (PR-1).

## Discussion

Host responses to biotic stress rely on the timely recognition of the pathogen and the efficient activation of a defense response that involves transcriptional reprogramming. The first stages of the interaction are decisive for the outcome of the disease, and therefore we focused on two early stages of *D. caulivora* infection (8 and 48 hpi), according to stem colonization process and induction of PR gene expression^2^. Comparative transcriptional profiles between Génesis 5601 and Williams detected more than 2.000 DEGs. From these, 46 % were commonly upregulated in both genotypes during *D. caulivora* infection, indicating overlapping responses. Interestingly, Génesis-8, showed 2.4 fold more upregulated DEGs than Williams-8 (1.028 versus 434 DEGs), while at 48 hpi the number of upregulated DEGs did not differ significantly between genotypes (1109 in Génesis-48 and 1205 in Williams-48). Comparative GO and pathway enrichment analysis resulted in similar terms and pathways in both infected genotypes, although Génesis-8 exhibited a greater number of genes in each category. Most enriched terms and pathways were related to plant defense, including defense response, response to oxidative stress and oxidant detoxification, phenylpropanoid and flavonoid biosynthesis, plant hormone signal transduction, plant-pathogen interaction, as well as protein phosphorylation and regulation of transcription. These finding suggest that a rapid recognition of the pathogen and activation of defense mechanism could lead to *D. caulivora* resistance in Génesis 5601.

The interplay between pathogen perception and defense activation have a profound effect on plant resistance. Our results revealed that 15 PRRs exhibited higher transcript levels in untreated Génesis 5601 compared to untreated Williams, including LRR-STKs, RLK and RLPs. This number increased to 59 upregulated PRRs in Génesis-8 versus 11 in William-8, including *RLKs, LRR-RLKs, LecRLKs, RLPs* and *CRKs*. Thus, basal PRRs expression levels as well as a rapid induction of PRRs upon *D. caulivora* inoculation might play important roles in PAMPs recognition and PTI activation. In accordance with these results, narrow-leafed lupin RLKs, LRR-RLKs, LecRLKs and RLP encoding genes were earlier induced in resistant compared to susceptible genotypes in response to *D. toxica*^23^. Similarly, several PRR such as CRKs and LRR-RLKs were significantly higher expressed in resistant compared to susceptible soybean plants in response to *Phakopsora pachyrhizi*^24^, soybean mosaic virus (SMV)^25^, and soybean cyst nematode^26^. Moreover, RLKs and RLPs are candidate resistance genes against the fungus *Phialophora gregata* (sin. *Cadophora gregata*)^27^.

Suppression of PTI by pathogen effectors activates ETI through the action of NLR^10^. The analysis of *D. caulivora* genome showed the presence of 133 secreted effector candidates, and several were induced during soybean colonization, suggesting that they could interfere with soybean defense^16^. Here, we show that seven *NLR* genes exhibited higher expression levels in untreated Génesis 5601 versus Williams. Similarly, seven and only one NLR were upregulated in Génesis-8 and Williams-8, respectively. This could lead to earlier activation of downstream immune events in Génesis 5601. Soybean NLR genes co-localize with disease-resistance quantitative-trait loci (QTL)^28^, supporting their role in plant resistance against different pathogens. NLR were identified as candidate resistant genes for several soybean diseases caused by bacterial, fungal, oomycete and virus^29-36^. Some of these NLR genes have been functionally validated^31,37,38^. Furthermore, genomic regions identified by genome-wide association studies (GWAS) and related to resistance against several soybean diseases were enriched in LRR-RLK and NLR^39^.

MAPKs, MAPKKs, MAPKKKs and STKs activate downstream signaling after pathogen recognition^40^. *D. caulivora* inoculation increased expression of genes encoding these type of kinases in both cultivars. Several *MAPKKKs* were only upregulated in Génesis-8 and STKs were differentially expressed in Génesis-8 and Williams-8. Interestingly, STKs are candidate genes for resistance to soybean mosaic virus and several of them were higher expressed in resistant compared to susceptible plants^25^.

WRKY, AP2/ERF, MYB, bHLH, NAC, GRAS, and other TFs play important roles in defense responses to biotic and abiotic stress^41-43^. Our study found that 219 of these TFs were upregulated during *D. caulivora* colonization. Génesis-8 showed significantly more upregulated *ERFs, MYB* and *bHLH* compared to Williams-8, which could trigger rapid induction of target genes involved in plant immunity. Similarly, higher expression levels of *bHLH* were observed in *P. sojae* resistant compared to susceptible soybean cultivars^44^. GmMYBs and GmNAC regulate biosynthesis of flavonoids leading to the production of phytoalexins that increase resistance against pathogen^45-47^. Moreover, GmERF5- and GmERF113-overexpressing soybean plants showed enhanced resistance to *P. sojae* and positively regulated expression of PR-10 and PR-1^48,49^.

A high number of PR proteins were induced during *D. caulivora* colonization in both cultivars, indicating their significant role during soybean defense against this fungal pathogen. Likewise, PR-1, β-1,3-glucanase (PR-2), chitinases (PR-3 and PR-4), PR-10 and defensin were induced in a susceptible cultivar infected with *D. aspalathi*^13^. Expression profiles highlighted β-1,3-glucanase, peroxidases class III (PR-9) and PR-10 function in the early resistant response of Génesis 5601. Similarly, peroxidases encoding genes were major upregulated DEGs in a resistant genotype of narrow-leafed lupin infected with *D. toxica*^23^. As observed for other pathogens, these enzymes could protect plant cells against *D. caulivora* by degrading fungal cell wall polysaccharides (β-1,3-glucanases), and probably by inhibiting hyphal growth through RNase activity (PR-10)^50,51^. Peroxidases could increase soybean defenses by reinforcing plant cell walls, synthesis of phytoalexins, or participate in the metabolism of ROS as has been observed in other pathogen-infected plants^52^. In addition to peroxidases, we found increased expression of other oxidative stress related genes encoding glutathione S-transferase, thioredoxin, ferredoxin oxidoreductases in Génesis-8, supporting the importance of cellular homeostasis to maintain a redox balance in soybean tissues to resist further infection. ROS accumulation can lead to a hypersensitive response (HR), a type of programmed cell death (PCD) that restricts the pathogen to the site of infection^53^. Interestingly, the induced expression of genes encoding BAG cochaperones in Génesis, suggest a possible involvement of PCD in *D. caulivora* resistance. These type of proteins trigger autophagy in the host to limit fungal colonization and confer resistance to fungal^54^.

Plants respond to pathogen infection by activating the phenylpropanoid pathway^14,55^. Here, we show that a high proportion of genes of this pathway are induced upon *D. caulivora* infection, indicating that reinforcement of the cell wall through lignin and phenolic compounds, synthesis of antimicrobial compounds such as flavonoids, isoflavonoids, coumarins and lignans are important defense mechanisms against this fungal pathogen. Interestingly, a significant number of upregulated DEGs were only present in Génesis-8 and not in Williams-8, suggesting that some of the metabolites produced by the phenylpropanoid pathway could be involved in resistant mechanisms against *D. caulivora*. Phytoalexin production in response to pathogens is regulated by the enzymes PAL, CHS, and chalcone isomerase (CHI), among others^14^. The number of upregulated DEGs encoding these enzymes were significantly higher in Génesis-8 compared to Williams-8. Interestingly, phenylpropanoids such as isoflavones daidzein, genistein and glyceollins are produced in soybean resistant plants after treatment with *D. aspalathi* elicitors^56^. Moreover, GmMYB29 regulates isoflavonoid biosynthesis in soybean through the activation of isoflavone synthase and CHS encoding genes^45^. Functional analysis demonstrated that GmMYB29A2 is essential for accumulation of glyceollin I and expression of *P. sojae* resistance^47^. Likewise, R2R3-MYB involved in lignin synthesis and genes responsive to chitin were significantly induced in *P. pachyrhizi* resistant genotypes^46^. Furthermore, activation of cell wall reinforcement by incorporation of phenolic compounds was previously observed in *D. caulivora* infected tissues^2^. Thus, these findings suggest that several metabolites of the phenylpropanoid could be produced during *D. caulivora* colonization, although further studies are needed to decipher the involvement of these metabolites in soybean resistant responses against *D. caulivora*.

Plant hormones conform a complex network that regulate plant resistance against pathogens^57^. Soybean plants activate hormonal pathways during *D. cualivora* infection in both genotypes. The significantly higher number of upregulated DEGs involved in IAA, ET, CK and JA pathways in Génesis-8 compared to Williams-8 suggests that these hormones could be involved in resistance responses against *D. caulivora*. Interestingly, induction of ACS and ACO genes involved in ET synthesis have been associated with resistance of soybean plants to vascular disease caused by *Fusarium virguliforme*^58^. In addition, genes encoding enzymes involved in the production of JA, LOXs and OPR, were faster induced in Génesis compared to Williams. Consistently, expression levels of LOXs increased in resistant genotypes of narrow-leafed lupin in response to *Diaporthe toxica*^23^. Oxylipins produced by the LOX pathway play different roles during defense responses against biotic stress through antimicrobial activities, contribution to HR, and production of signaling molecules such as JA and related compounds that lead to induction of genes with multiple roles in defense^15,59^.

Small HSP are chaperones that play important roles in immunity by protecting cells from stress-induced protein aggregation and misfolding^60^. Remarkably, sHSPs encoding genes were only upregulated in Génesis and generally at 8 hpi, suggesting their involvement in the early stages of plant resistance. HSPs are involved in stability and accumulation of PRRs and NLRs and further defense signaling^60,61^. In *Nicotiana tabacum*, a sHSP was shown to be involved in disease resistance against *Ralstonia solanacearum*^62^. Furthermore, GmHsp22.4 was highly induced in a nematode resistant soybean genotype and its overexpression in Arabidopsis plants renders lower nematode multiplication^63^. Interestingly, some pathogen effectors interact with sHSPs to suppress chaperone activity and promote virulence^64^, highlighting the role of these plant chaperones in disease resistance.

## Conclusions

This comparative transcriptomic study between contrasting soybean genotypes revealed that the observed resistance of Génesis 5601 to *D. caulivora* infection, is likely based on a rapid recognition of the pathogen and induction of genes related to plant immunity. Soybean defense activation to *D. caulivora* involves perception through PRR and NLR, as well as TFs, PRs, biosynthesis of phenylpropanoid derived metabolites, hormones, sHSPs and genes with different roles in defense. Future studies involving functional characterization of soybean candidate genes and target genes of *D. caulivora* effectors will contribute to a valuable comprehension of soybean-*Diaporthe* interactions. These findings provide novel insights into soybean defense molecular mechanisms used to control this pathogen, and establish a foundation for improving resistance in breeding programs.

## Methods

### Plant materials and *D. caulivora* inoculation

A *D. caulivora* isolate (strain D57), collected from canker lesions of soybean in Uruguay during 2015 was used in this study^2^. Two soybean genotypes were used for all plant assays: SSC-susceptible Williams PI548631 obtained from USDA ARS Soybean germplasm collection (seed source 13U-9280, order 253444, 2014), and the SSC-resistant Génesis-5601 from the Instituto Nacional de Investigación Agropecuaria (INIA) breeding program (Stewart S, personal communication). Three soybean seeds of each genotype were individually planted in a 10-cm-diameter pot filled with a mix of soil and vermiculite at a rate of 3:1. Soybean seedlings were grown in a growth room under a 16 h light/8 h dark photoperiod regime at 24°C. For all experiments, 3-week-old plants at V2 were used. *D. caulivora* D57 isolate was inoculated using the stem wounding method where an agar plug containing mycelium was applied to the wounded stem^2^. As a control an agar plug without mycelium was used. All experiments were performed with accordance to relevant regulations and guidelines.

Development of SSC symptoms were compared between both soybean genotypes. Ten plants were used per treatment and the experiment was repeated three times. Lesion length (mm) and disease severity (Scale 1-7), was determined at various time points (3, 5, 7, 11, and 14 dpi]. Disease severity index and area under disease progress curve (AUDPC) was calculated according to Mena et al.^2^. Differences between treatments were determined by non-parametric Kruskal–Wallis and Mann– Whitney tests using SPSS Statistics v. 21.0. The significance level for data used was p < 0.01.

### Quantitative PCR

After soybean stem inoculation with *D. caulivora*, fungal DNA was quantified at 8, 24, 48, 72, and 96 hpi. Three plants per treatment were used as biological replicates and samples were frozen in liquid nitrogen. DNA extraction, quantitative PCR (qPCR), estimation of pathogen biomass and the amount of soybean DNA was performed according to Mena et al.^2^. For qPCR, primers designed for the elongation factor gene of soybean and the β-tubulin gene of *D. caulivora* were used. Results were expressed as ng of *D. caulivora*/ng of soybean tissue. Student’s t-test was applied to all qPCR data, and values of p ≤ 0.01 were considered statistically significant.

### RNA extraction, cDNA library preparation and sequencing

Samples of both genotypes were taken from untreated plants, and at 8 and 48 hpi inoculated with plugs containing *D. caulivora* mycelium and their respective controls (plugs without mycelium) were used for transcriptomic analysis. Total RNA was extracted and purified from 100 mg soybean stems, 10 mm above the inoculation point of each sample with TRIzol reagent (Invitrogen, Carlsbad, CA, USA) and Invitrogen PureLink RNA Extraction Mini kit (Invitrogen, USA), including an on-column digestion with RNase-Free DNase I, were used according to the manufacturer’s instructions. RNA quality was checked by running samples on 1.2% formaldehyde agarose gel. RNA concentration was measured using a NanoDrop 2000c (Thermo Scientific, Wilmington, USA). RNA quality control, library preparation, and sequencing were performed at Macrogen Inc. (Seoul, Korea). Three biological replicates were included per treatment. Libraries for each biological replicate were prepared for paired-end sequencing by TruSeq Stranded Total RNA LT Sample Prep Kit (Plant) with 1 g input RNA, following the TruSeq Stranded Total RNA Sample Prep Guide, Part # 15,031,048 Rev. E. Sequencing was performed on Illumina platform (Illumina, CA, USA) to generate paired-end 101 bp reads, obtaining 41.6 to 65.2 M reads per sample with Q20 > 98% and Q30 > 95%.

### Pre-processing of raw data, mapping of reads and annotation

RNA-seq processing steps were done through Galaxy platform (https://usegalaxy.org/)^65^. Raw reads quality was subjected to a quality control check using FastQC software ver. 0.11.2 (http://www.bioinformatics.babraham.ac.uk/projects/fastqc/). Sequences were trimmed, and the adapters removed using Trimmomatic Version 0.38.0 software^66^. Additionally, to the default options, the following parameters were adjusted: adapter sequence TruSeq3 (paired-ended (PE), for MiSeq and HiSeq), always kept both PE reads, and SLIDINGWINDOW: 4:15 HEADCROP: 13 MINLEN:50. Trimmed reads were mapped to reference genome of Glycine max Gmax_275_Wm82.a2.v1.fa^67^ as the reference genome file and Gmax_275_Wm82.a2.v1.gene.gff3 as a reference file for annotation gene models from Phytozome (https://phytozome.jgi.doe.gov/pz/portal.html) using Hisat2 software^68^. The BAM files were obtained with Samtools View software ver.1.9 and then sorted by name with Samtools Sort software ver. 2.0.3^69^, for further analysis. Reads were counted using FeatureCounts software ver. 1.6.4^70^. Additionally, to default options, parameters were adjusted for: count fragments instead of reads, allow read to map to multiple features, count multi-mapping reads/fragments and use reference sequence file *Glycine max* v4.2.

### Transcript expression and functional analysis

Cluster analysis of replicates from each time point and control samples were performed by Principal Component Analysis (PCA) using log2 fold changes for all datasets. Differential expression analyses were performed using EdgeR software ver. 3.24.1^71^, with p-value adjusted method of Benjamini and Hochberg adjusted threshold 0.05^72^, and Minimum log2 Fold Change 2. Counts were normalized to counts per million (cpm) with the TMM method and low expressed genes filtered for count values ≥ 3 in all samples. In this study, a false discovery rate (FDR) ≤ 0.05 was used to determine significant differentially expressed genes (DEGs) between *D. caulivora* inoculated plants and mock; and expression values were represented by log2 ratio. Heat maps were generated using the Heatmapper server (www.heatmapper.ca/expression). Hierarchical clustering analysis of expressed genes were performed on log2 Fold-Change expression values using the “hclust” tool from R package “stats” ver. 3.6.0. To visualize the obtained data, heatmap plots were performed using the “heatmap.2” tool from R package “gplots” ver. 3.1.0.

Gene ontology (GO) and functional annotations were assigned with the Blast2GO, through Omicbox software (https://www.biobam.com/omicsbox)^73^. Gene models were compared with several databases (NCBI nonredundant protein database, GO, and InterpoScan) with BlastP finding single hit at an e-value threshold of e-value ≤ 1.0E-3 using taxIds for Viridiplantae. InterproScan analysis was used to identify domains in the genome^74^ DEG functional enrichment analysis was performed using OmicBox software. GO terms with a FDR ≤0.05 were considered for the analysis. DEGs of each dataset were divided into upregulated and downregulated subsets. Kyoto Encyclopedia of Genes and Genomes (KEGG) enrichment analysis of all DEGs were obtained through Omicbox software for *D. caulivora* inoculated vs. control samples in both soybean genotypes. All heat maps were generated using the Heatmapper server (www.heatmapper.ca/expression).

### Data availability

RNA sequencing data were deposited at the National Center for Biotechnology Information (NCBI) in the Sequence Read Archive (SRA) under the PRJNA878492 Bioproject accession.

## Acknowledgements

Authors thank Ricardo Larraya for technical assistance. This work was supported by “Agencia Nacional de Investigación e Innovación (ANII) (graduate fellowship)” Uruguay, “Programa de Desarrollo de las Ciencias Básicas (PEDECIBA)” Uruguay, and “Programa Grupo de I+D Comisión Sectorial de Investigación Científica, Universidad de la República”, Uruguay.

## Author Contributions

EM performed all the experiments, interpreted the data, contributed to discussions, and helped to write the manuscript. GR performed the hierarchical clustering analysis. SS contributed to the selection of the soybean genotypes and discussions. MM contributed to discussions and helped to write the manuscript. IP designed and supervised the study, interpreted the data, and wrote the manuscript. All authors read and approved the final manuscript.

## Supporting information

All code/additional information needed for the full reproduction of this study will be made available upon acceptance of this manuscript.

## Additional information

**Supplementary Figure 1:** Two-dimensional scatterplot of the principal component analyses (PCA) for soybeans where distances approximate the typical log2 fold changes between the samples. Colored dots denote each biological replicate.

**Supplementary Table S1:** Summary of mapped reads of Williams and Génesis 5601 RNA-Seq libraries. 1-3 indicate the three biological replicates in control stems and during *D. caulivora* stem infection at the indicated time points.

**Supplementary Table S2**. List of soybean differentially expressed genes (DEGs). Comparisons were performed between untreated Génesis 5601 versus untreated Williams (G-W), and *D. caulivora* inoculated versus control plants of both genotypes at the indicated time points (Gi8-G8; Gi48-G48; Wi8-W8; Wi48-W48).

**Supplementary Table S3:** List of all DEGs in soybean plants inoculated with *D. caulivora* obtained by hierarchical clustering.

**Supplementary Table S4:** List of DEGs encoding for proteins with role in perception and signaling during *D. caulivora* infection in both genotypes at 8 and 48 hpi.

**Supplementary Table S5:** List of DEGs encoding for proteins involved in transcription during *D. caulivora* infection in both genotypes at 8 and 48 hpi.

**Supplementary Table S6:** List of DEGs encoding for pathogenesis related proteins during *D. caulivora* infection in both genotypes at 8 and 48 hpi.

**Supplementary Table S7:** List of DEGs encoding for proteins involved in phenylpropanoids and flavonoids pathways during *D. caulivora* infection in both genotypes at 8 and 48 hpi.

**Supplementary Table S8:** List of DEGs involved in hormone signaling during *D. caulivora* infection in both genotypes at 8 and 48 hpi.

